# PathPCNet: Pathway Principal Component-Based Interpretable Framework for Drug Sensitivity Prediction

**DOI:** 10.1101/2025.08.20.668802

**Authors:** Bikhyat Adhikari, Masrur Sobhan, Ananda Sutradhar, Giri Narasimhan, Ananda Mohan Mondal

## Abstract

Precision medicine aims to identify significant biomarkers and effective drugs tailored to individual genomic profiles, thereby enabling personalized treatment strategies. Drug efficacy is often attributed to drug response, commonly measured as the concentration of a drug required to inhibit a biological activity. In contrast, drug sensitivity reflects how strongly a tumor responds to a drug, where a lower effective dose indicates higher sensitivity. Machine learning-based drug response prediction has the potential to accelerate biomarker discovery and facilitate the development of more effective therapeutics.

In this study, we present *PathPCNet*, a novel interpretable deep learning framework that integrates multi-omics data (copy number variation, mutation, and RNA sequencing) fused with biological pathways, drug molecular structure, and Principal Component Analysis for drug response prediction. Our model achieves a Pearson correlation coefficient of 0.941 and an R-squared of 0.885, outperforming the existing pathway-based approaches.

We employ SHAP-based model interpretation to quantify the contributions of omics and drug features, uncovering key pathways and gene-drug interactions associated with resistance mechanisms. These results demonstrate the utility of integrative deep learning models not only for accurate prediction but also for generating biologically meaningful insights, which can advance drug discovery and precision oncology. In addition, the framework also facilitates the identification of important pathways, genes, and atomic attributes of drugs related to drug sensitivity and different cancer types.

## 1 Introduction

The fundamental objective of precision medicine is to design the right treatment strategy for a patient based on the individual’s genetic profile [1], [2]. It involves identifying the significant biomarkers for tumor responses, as well as the efficient drugs [3]. Efficacy of drugs, often quantified as drug response, is commonly represented by half-maximal inhibitory concentration (IC50); a lower IC50 indicates a higher potency, meaning the drug is effective at a lower concentration [4], [5]. In contrast, drug sensitivity describes how strongly or weakly a biological system responds to a drug. A sensitive system shows a strong response at a low dose (low IC50), while a resistant system shows a weak or no response even at high doses (high IC50 or no inhibition). Therefore, accurately predicting drug response is crucial in precision medicine, as it allows tailoring therapies based on individual molecular profiles. Moreover, drug response modeling can aid in biomarker discovery and development of more effective therapeutics by linking molecular features with tumor-specific drug efficacy [6].

With the rise of large-scale multi-omics datasets, such as Cancer Cell Line Encyclopedia (CCLE) [7], Genomics of Drug Sensitivity in Cancer (GDSC) [8] and The Cancer Genome Atlas (TCGA) [9], machine learning and deep learning approaches have emerged as powerful tools for drug response studies. Despite their success, a vast majority of existing solutions offer limited interpretability and fail to elucidate underlying biological mechanisms driving their predictions [10]. To bridge this gap, several recent studies [11]–[15] have attempted to develop biologically informed and interpretable ML frameworks. However, the feature selection and extraction methods most of these studies apply rely solely on statistical or machine learning heuristics, which may not possess proper biological relevance. In response, several recent works have incorporated biological pathways to enhance interpretability and biological relevance in precision medicine [16]–[19].

For example, Tang and Gottlieb [16] implemented an interpretable deep learning framework for drug response prediction based on pathway enrichment scores. The authors used Lundberg and Lee’s SHAP [20] framework – a widely adopted model-agnostic explainable method – to identify the feature contributions. However, a major drawback of enrichment score is that it is not possible to go back to the original features (genes) from the pathway enrichment score. Thus, their explanation is limited to only the pathway features.

In this study, we present **PathPCNet**, a novel, interpretable deep learning framework that leverages biological pathways, multi-omics data, and Principal Component Analysis (PCA) for drug response prediction. Instead of using raw gene-level features – which are often high-dimensional and noisy – we project cell line features onto pathway-level principal components (Pathway PCs) derived from curated gene sets in Pathway Interaction Database (PID) [21]. Based on the experiments conducted by Eckhart et al. [3], PCA is a top-performing dimensionality reduction technique for drug response prediction. The pathway PCA-based transformation not only reduces dimensionality but also enhances biological interpretability by preserving relevant variations at the pathway level. Furthermore, we employ SHAP to interpret the model and to identify significant features that influence the tumor response. Additionally, we back-project the SHAP scores from pathways to the original features using PCA loadings to identify the most significant genes. To the best of our knowledge, this is the first work to apply PCA on pathway-based multi-omics data for drug response prediction.

## 2 Materials and Methods

### A. Data Overview

Table I shows the data overview. For this study, we used three types of omics data (Copy Number Variation, mutation, and RNA seq), drug response data, Morgan fingerprint of drugs, and pathway data. We used release 8.5 of Genomics of Drug Sensitivity in Cancer (GDSC2) [22] dataset. We retrieved omics data from Cell Model Passports repository [23]. The drug dataset downloaded from the GDSC portal comprised 297 unique compounds (by drug ID). GDSC2 drug response data contains log-normalized IC50 values for 969 unique cell lines and 295 unique drugs. Additionally, we used PubChem database [24] to obtain SMILES (Simplified Molecular-Input Line-Entry System) [25] strings for drugs. Further, C2 curated canonical pathway and genes from Pathway Interaction Database (PID) [21] were obtained through Molecular Signatures Database (MSigDB) [26], [27]. The pathway data includes 2,534 unique genes spanning 196 pathways.

**TABLE I:** Overview of datasets. The final data used in this study are indicated by bold-face.

| Omics Data Overview |  |  |  |  |  |  |  |
| --- | --- | --- | --- | --- | --- | --- | --- |
| Data Type | Number of Cell lines |  |  |  | Number of Genes |  |  |
|  | Original | Sanger Source | GDSC2 | Filtered (for PCA) | Original | Pathway | Filtered (for PCA) |
| CNV | 1,253 | 1,033 | 960 | **409** | 18,348 | 2,513 | **2,509** |
| Mutation | 1,435 | 1,215 | 964 | **409** | 23,189 | 2,522 | **2,522** |
| RNA Seq | 1,431 | 442 | 412 | **409** | 37,600 | 2,529 | **2,529** |
| Drug Response Overview |  |  |  |  |  |  |  |
| Drug Response | Number of Cell lines |  | Number of Drugs |  | Number of Drug Response |  | Cancer Types |
|  | Original | Filtered | Original | Filtered | Original | Filtered |  |
| Drug Response | 969 | **409** | 295 | **182** | 242,036 | **67,279** | 29 |
| Morgan Fingerprint: 256 Bits Representation of 182 Drugs |  |  |  |  |  |  |  |
| Pathway data: 196 pathways with 2,534 unique genes |  |  |  |  |  |  |  |

### B. Preprocessing Omics Data

Copy Number Variation (CNV) data was available in long format, where each row is identified by a cell line and gene pair. We filtered pathway genes for Sanger cell lines and applied the GISTIC2 (Genomic Identification of Significant Targets in Cancer version 2.0) threshold [28] to the copy number categories as follows: −2 (Deletion), −1 (Loss), 0 (Neutral), 1 (Gain), and 2 (Amplification). The processed data was transformed into a matrix with discrete values. The transformed matrix had 2,513 pathway genes, of which 102 genes had some missing values. Among these 102 genes, 98 genes had missing CNVs for less than 15% of the cell lines; we imputed these with a zero (indicating Neutral), and discarded the other four genes (SRY, PPP2R3B, IL3RA, and CSF2RA) as they had over 50% of the values missing.

Mutation data was available in the form of Variant Allele Frequency (VAF) in long format. We filtered this data to retain only entries derived from the Sanger source and limited gene selection to those present in curated biological pathways, and transformed it to a matrix of cell lines and genes. After transformation, the missing values in the resulting matrix were imputed with zeros, indicating the absence of mutation.

RNA sequencing expression data, represented as Transcript Per Million (TPM) values, was also in long format. Similar to other omics data, we retained only Sanger-derived entries and filtered genes to those associated with pathways. The data was reshaped into a two-dimensional matrix, and a base-2 log transformation was applied.

After aligning all datasets, we identified 409 cell lines common across drug response, CNV, mutation, and gene expression data. Next, we normalized the CNV and gene expression data in the range of [0,1] to bring them all to the same scale. The final datasets consist of 2,509 genes in CNV, 2,522 genes in mutation, and 2,529 genes in expression data, each for 409 cell lines, as shown in Table I in bold-face.

### C. Preprocessing Drug Data

Drugs with missing or multiple PubChem IDs and those missing from the PubChem database [24] were excluded. For the remaining drugs, we used RDKit [29] to generate Morgan fingerprints with a dimension of 256 bits from drug SMILES retrieved using PubChemPy API and the PubChem ID of drugs. These fingerprints are equivalent to Extended Connectivity Fingerprints (ECFPs) [30], a widely used molecular representation in cheminformatics. The final drug dataset comprises 182 unique compounds (by drug ID), each represented by 256-bit Morgan fingerprints, as shown in Table I.

### D. Processed Data

The final processed data contains 67,279 drug response values for 409 unique cell lines (314 cell lines with known cancer type, 90 cell lines with unclassified label, and 5 cell lines with missing cancer type information) and 182 unique drugs. It should be noted that for this study, missing cancer type information has no impact. Not every drug has the response value for all cell lines. The number of cell lines per drug ranges from 117 to 409, covering 29 different types of cancer from the GDSC2 cell lines. Table I shows the summary of the data used in this study.

### E. Proposed Pipeline

Fig. 1 shows the overall workflow of PathPCNet for drug response prediction. We calculate the pathway-specific principal components for all omics data. Then, PC features are combined with the Morgan fingerprint of all the drugs along with the IC50 values for each cell line and drug pair to create the input data matrix with 67,279 rows for regression models.

**Fig. 1:**
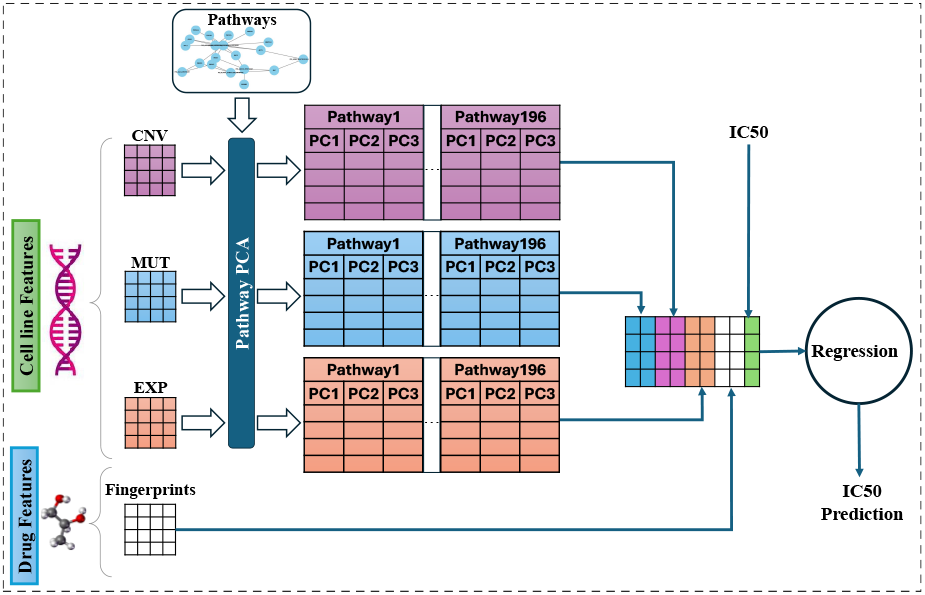
Workflow pipeline for PathPCNet. Processed omics data (CNV, MUT, EXP) are first projected to principal components (PCs) for individual pathways, then these PCs and drug features are used as the input matrix for the regression task. CNV: Copy Number Variation; MUT: Mutation; EXP: RNA Seq Expression; PCA: Principal Component Analysis.

### F. Principal Component Analysis and Model Selection

We calculated the first four principal components for all 196 pathways for the three omics (CNV, mutation, and expression) data for 409 common cell lines from the preprocessed data. This process resulted in 196 *×* 4 = 784 pathway features for each of the three omics datasets.

Next, we trained six different regression models: XG-Boost [31], LightGBM [32], Extra Trees [33], Ridge [34], Random Forest [35], and a neural network. These models were trained using one to four principal components and all drug features, with ten-fold cross-validation, implemented using Python libraries [36], [37]. Based on the average values of evaluation metrics, as shown in Fig. 2, the neural network provides the best performance. The neural network is a multi-layer perceptron (MLP), adapted from PathDSP [16], consisting of an input layer (matching the combined size of PCA features and the 256-bit drug fingerprint), followed by four hidden layers with 1000, 800, 500, and 100 neurons, each using ELU activations. The output layer is a single linear unit predicting IC50. Additionally, increasing the number of principal components does not improve IC50 prediction performance. Hence, we selected neural network as our regression method and used only first principal component (PC1) for further analysis. PC1 alone captures 17% of data variance on average.

**Fig. 2:**
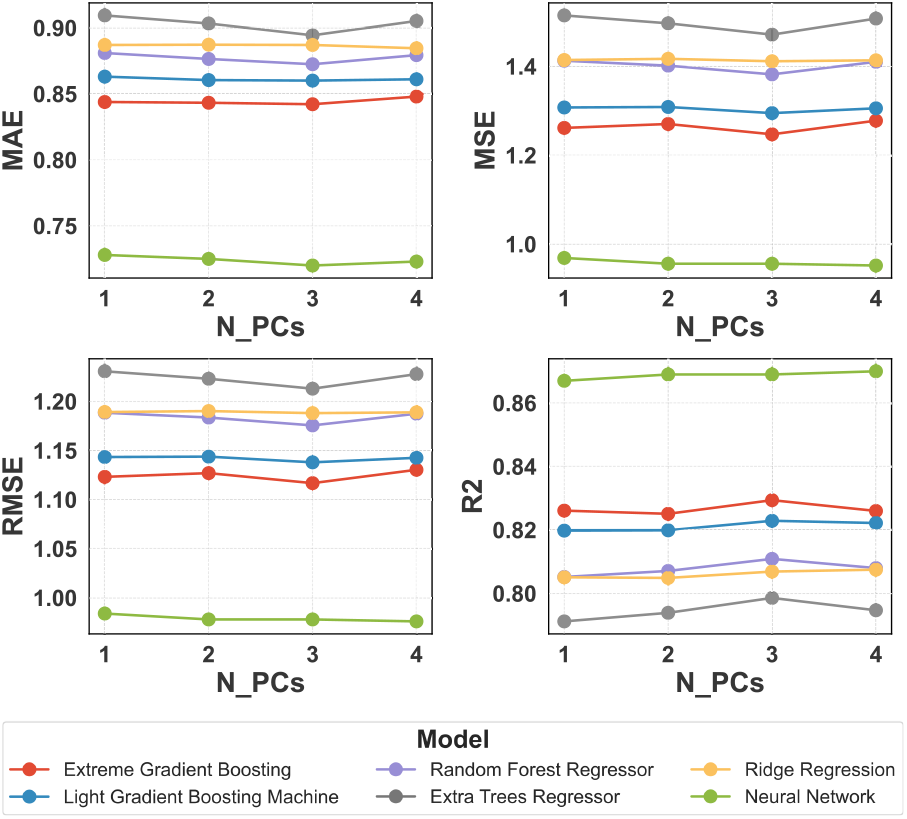
Regression evaluation metrics (average from ten-fold cross validations) across six different models for varying numbers of principal components (N_PCs). MAE: Mean Absolute Error; MSE: Mean Squared Error; RMSE: Root Mean Squared Error; R^2^: Coefficient of Determination.

### G. Hyperparameter Tuning

Table II summarizes the hyperparameter values explored during the neural network tuning. The tuning was performed based on RMSE using an 80:20 train–test split. Within the training set, 10% of the data was further set aside for validation. The hyperparameter combination that resulted in the lowest validation RMSE of **0.92** was selected as optimal and is highlighted in Table II. These optimal values were used in all subsequent analyses and for reporting the final results.

**TABLE II:** Hyperparameter tuning setup. Optimal hyperparameters are in bold-face.

| Hyperparameter | Values |
| --- | --- |
| Learning Rate | $[1 \times 10^{-3}, 5 \times 10^{-4}, \mathbf{1 \times 10^{-4}}, 5 \times 10^{-5}]$ |
| Dropout Rate | $[0.1, 0.2, \mathbf{0.3}, 0.4, 0.5]$ |
| Batch Size | $[8, 12, 16, \mathbf{20}, 24, 28]$ |
| Optimizer | adam, **adamw**, rmsprop, sgd (momentum=0.9) |

## III. Results

### A. Predictive Performance

Fig. 3 shows the predictive performance PathPCNet. It accurately predicts the drug response by integrating multiomics data with biological pathways and drugs’ molecular structures.Thefinal regressionoutputof themodelhasMAE of0.677±0.005,PearsonCorrelationCoefficient (PCC)of 0.941±0.001, andR-squaredof0.885±0.001

**Fig. 3:**
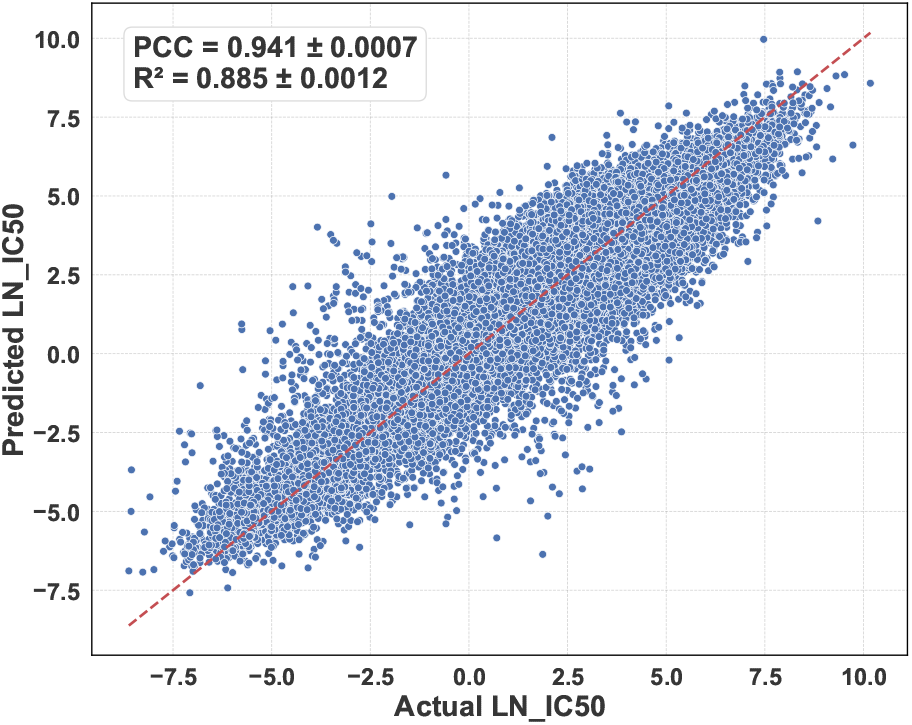
Performance of PathPCNet for drug response prediction. LN_IC50: Log Normalized IC50, PCC: Pearson Correlation Coefficient, *R*^2^: Coefficient of Determination.

### B. Significant Pathways

We applied SHAP [20] Deep Explainer on the trained neural network model to calculate the feature contribution for the model’s output. We took the average of absolute SHAP values to rank the feature contribution. Further, we calculated the distribution of the top 200 features: a snapshot of the distribution is presented in Table III.

**TABLE III:** Snapshot of top N feature distribution based on SHAP values.

| Top N Features | CNV | Mutation | Expression | Fingerprint |
| --- | --- | --- | --- | --- |
| 10 | 0 | 0 | 0 | 10 |
| 50 | 0 | 8 | 5 | 37 |
| 100 | 0 | 19 | 18 | 63 |
| 150 | 4 | 38 | 27 | 81 |
| 200 | 12 | 46 | 42 | 100 |

Next, we wanted to identify the most significant pathways based on the combined importance of all the omics data. Fig. 4 shows the top ten pathways based on the combined feature importance. Interestingly, the pathway ‘PID_PS1_PATHWAY‘ appears for all three omics data, which underscores its significance in the drug response. This pathway has 46 genes, out of which the genes ‘HDAC1‘ and ‘GSK3B‘ also appear in the target genes in the GDSC2 drug dataset. The second most significant pathway is ‘PID_TAP63_PATHWAY‘. Among 54 genes in this pathway, ‘MDM2‘, ‘PLK1‘, ‘SP1‘, ‘MDM2‘, and ‘EP300‘ appear in the target genes in the GDSC2 drug dataset.

**Fig. 4:**
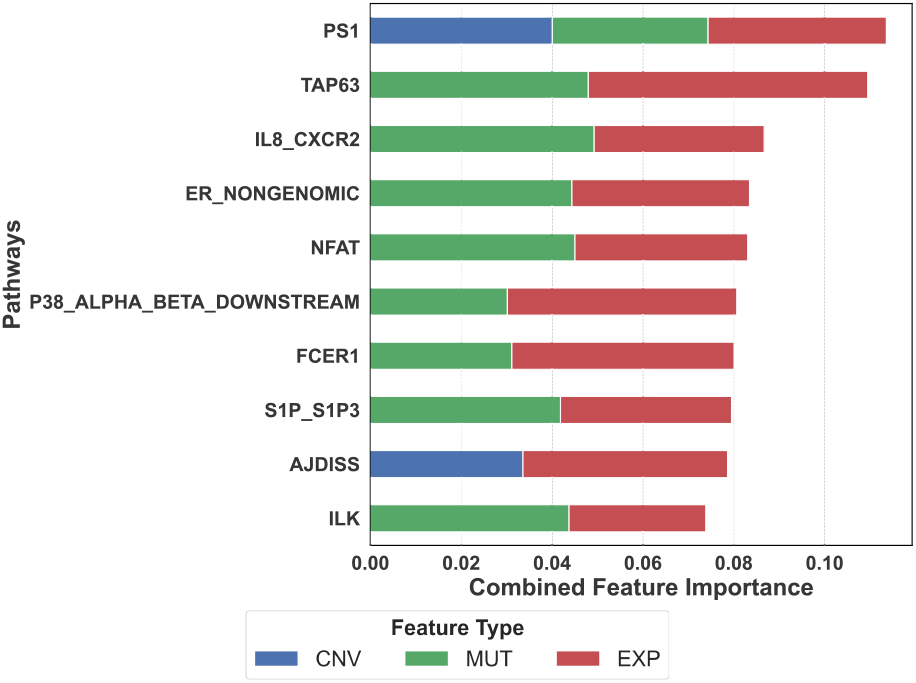
Top 10 pathway features based on average feature contribution of different omics data towards IC50 prediction. Y-axis labels are pathway names, e.g., PS1 is “PID_PS1_PATHWAY”.

### C. Significant Genes

We back-projected pathway principal component features’ SHAP scores to the original feature dimension (genes) to identify significant genes for all omics data. Since the same gene appears in multiple pathways and different omics, we calculated the absolute sum of feature contribution of all genes across different pathways in different omics data to rank the genes based on their significance. Fig. 5 shows the top ten genes based on the calculated feature contributions for the genes. Four genes (‘SRC‘, ‘RAC1‘, ‘EP300‘, ‘EGFR‘) among these top ten genes appear in the drug targets in GDSC2, suggesting that PathPCNet captures the significant genes for drug response. For further validation, we downloaded the drug gene interaction data from DGIdb [38] for all 182 drugs. Out of 552 common genes between the drug gene interaction and the pathway genes, **117** genes are among the top 200 genes we ranked. Such a high number of overlaps suggests that our proposed pipeline properly captures the significant biomarkers.

**Fig. 5:**
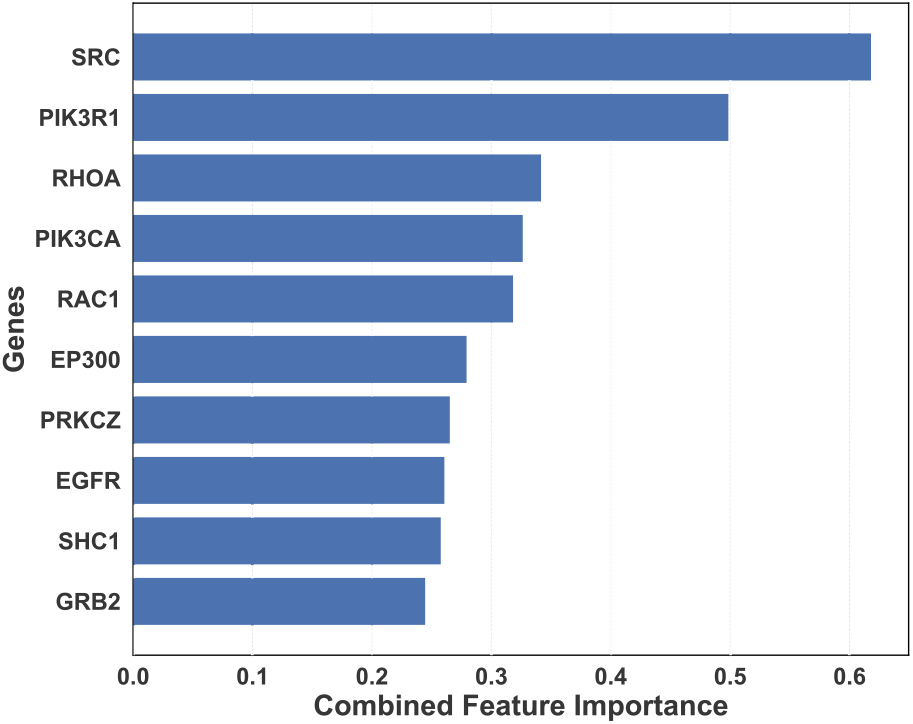
Top ten genes based on the combined values after back-projecting SHAP values using PCA loadings. As shown in Fig. 6, most atoms exhibit positive SHAP values, indicating that they contribute to higher predicted drug response values, i.e., greater resistance.

**Fig. 6:**
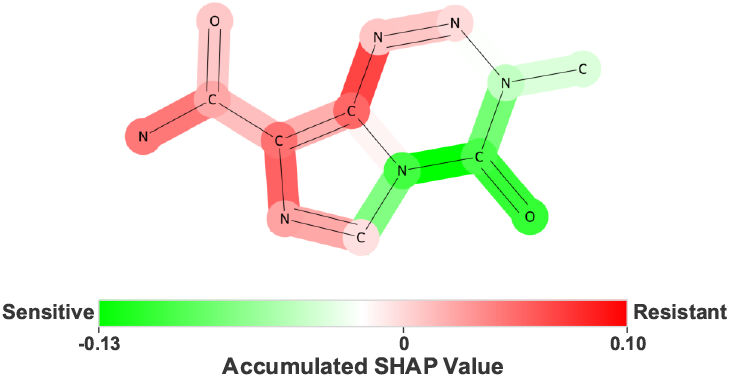
SHAP-based interpretation of atom contributions for drug response of Temozolomide for OVKATE cell line. The molecule is colored by accumulated SHAP values, with the colorbar indicating sensitivity to resistance. A darker red color indicates a higher contribution to making the drug resistant.

### D. Interpreting SHAP values for Drug Feature

We picked the drug response for the ‘OVKATE‘ cell line (https://www.cancerrxgene.org/cellline/OVKATE/1240199) and the ‘Temozolomide‘ drug to interpret the molecular basis of the model predictions. This particular drug response is very high, i.e., resistant. We back-projected the SHAP values from Morgan fingerprints to individual atoms in the SMILES of the drug. Specifically, we calculated atom-level contributions by summing the SHAP values of all fingerprint bits in which each atom participated. For each active bit, we identified its corresponding atomic environment using RDKit’s bitInfo and FindAtomEnvironmentOfRadiusN functions. The SHAP value of the bit was then equally distributed across all atoms involved in that substructure. This allowed us to visualize the accumulated SHAP importance over the molecular structure.

### E. Drug Specific Feature Importance

Next, we calculated the average of absolute SHAP values for each drug to identify the key pathway features from different omics that influence response for each of the drugs. We listed the top ten pathway features and the corresponding average SHAP scores for all 182 drugs. We calculated the frequency of these top ten pathway features to identify how many drugs and omics types share the same significant pathways, Fig. 7. The gene expression feature for the pathway ‘PID_CMYB_PATHWAY‘ appears in the top ten significant pathways for 169 out of 182 drugs. While further biological validation is required, the high frequency of ‘PID_CMYB_PATHWAY‘ among top predictive features underscores its importance in shaping drug response patterns. Moreover, SHAP scores for these significant pathways can be back-projected to the gene level to further study how significant genes would influence the drug responses.

**Fig. 7:**
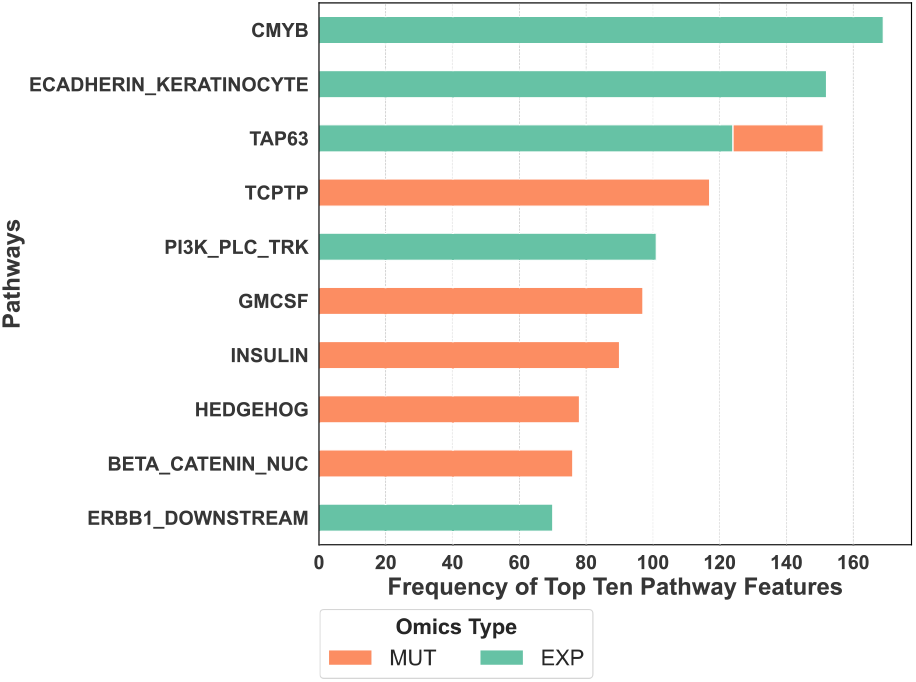
Top ten most frequent pathways based on the top ten most significant pathway features per drug. Y-axis labels are pathway names, e.g., CMYB is “PID_CMYB_PATHWAY”.

### F. Feature Importance Based on Cancer Types

We calculated the feature importance based on the cancer type labels of the cell lines following the same procedure as described for identifying feature importance based on the drug in Section III-E. Fig. 8 shows the top ten most significant pathways based on the cancer type of the cell lines. The gene expression feature for the pathway “PID_P38_ALPHA_BETA_DOWNSTREAM_PATHWAY” appears as the top ten most significant features for 11 different cancer type labels, which underscores its importance for different cancer types to study drug response. Additionally, PCA loadings can be used to identify the significant genes important to study drug response for different cancer types.

**Fig. 8:**
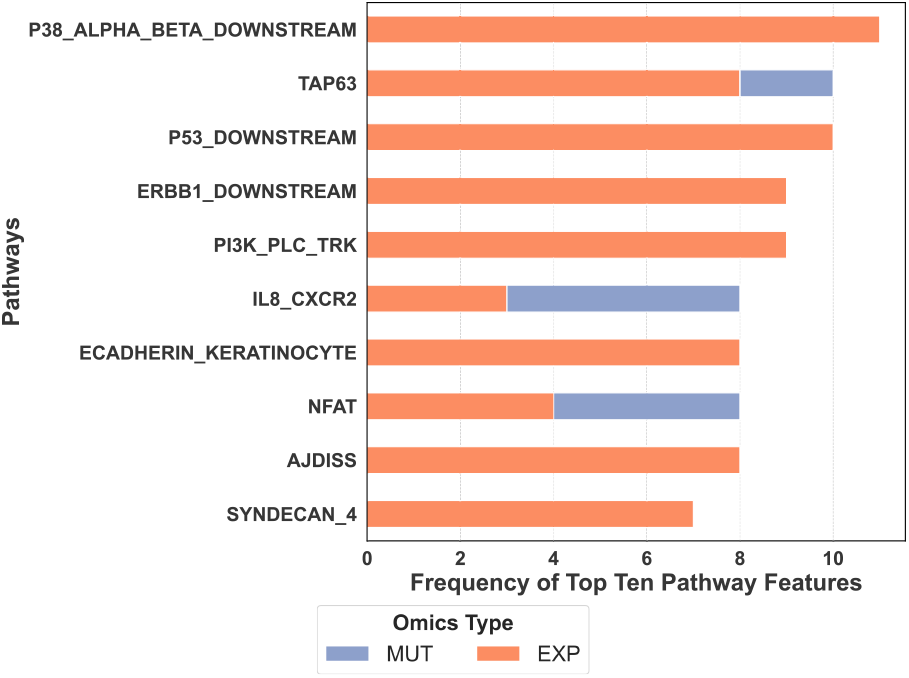
Top ten most frequent pathways based on the top ten most significant pathway features for each cancer type. Y-axis labels are pathway names, e.g., SYNDECAN_4 is “PID_SYNDECAN_4_PATHWAY”.

### G. Comparison with Other Methods

We compared the predictive performance of PathPCNet with two similar pathway-based drug response frameworks: PathDSP [16] and PASO [39] using our preprocessed data. Based on the average regression metrics from ten-fold cross validation presented in Table IV, our framework provides improved performance for drug response prediction, while providing the interpretation at the original feature dimension.

**TABLE IV:** Performance comparison with other pathway-based approaches (mean ± std)

| Model | MAE | RMSE | R<sup>2</sup> | PCC |
| --- | --- | --- | --- | --- |
| PathPCNet | 0.67 ± 0.005 | 0.91 ± 0.006 | 0.88 ± 0.001 | 0.94 ± 0.001 |
| PASO | 0.70 ± 0.005 | 0.95 ± 0.010 | 0.86 ± 0.004 | 0.93 ± 0.002 |
| PathDSP | 0.72 ± 0.011 | 0.99 ± 0.019 | 0.85 ± 0.005 | 0.92 ± 0.003 |

### H. Ablation Study

We evaluated the framework against the different modality, as shown in Table VI. Based on our experiment, using all three omics data provides the best predictive performance. Additionally, we evaluated our model with gene features (Table V). Our pathway principal component-based framework provides better predictive performance compared to raw gene features.

**TABLE V:** Performance comparison between first principal component features and raw gene features (mean ± std)

| Feature Type | MAE | RMSE | R<sup>2</sup> | PCC |
| --- | --- | --- | --- | --- |
| Pathway PC1 | **0.67 ± 0.005** | **0.91 ± 0.006** | **0.88 ± 0.001** | **0.94 ± 0.001** |
| Raw Gene | 0.86 ± 0.018 | 1.14 ± 0.026 | 0.82 ± 0.008 | 0.91 ± 0.004 |

**TABLE VI:** Evaluation of Multi-Omics Modality Contributions to Model Performance (mean ± std)

| Omics Modality | MAE | RMSE | R<sup>2</sup> | PCC |
| --- | --- | --- | --- | --- |
| CNV, EXP, MUT | **0.67 ± 0.005** | **0.91 ± 0.006** | **0.88 ± 0.001** | **0.94 ± 0.001** |
| CNV, EXP | 0.69 ± 0.008 | 0.94 ± 0.012 | 0.88 ± 0.003 | 0.94 ± 0.001 |
| EXP, MUT | 0.68 ± 0.006 | 0.93 ± 0.009 | 0.88 ± 0.002 | 0.94 ± 0.001 |
| CNV, MUT | 0.70 ± 0.008 | 0.95 ± 0.011 | 0.88 ± 0.003 | 0.94 ± 0.002 |
| CNV | 0.74 ± 0.008 | 1.00 ± 0.014 | 0.86 ± 0.004 | 0.93 ± 0.002 |
| EXP | 0.70 ± 0.007 | 0.96 ± 0.014 | 0.87 ± 0.004 | 0.94 ± 0.002 |
| MUT | 0.72 ± 0.008 | 0.98 ± 0.014 | 0.87 ± 0.003 | 0.93 ± 0.002 |

### I. Statistical and Literature Validation

We performed pathway enrichment analysis on top 200 SHAP-derived genes using hypergeometric test with Benjamini-Hochberg FDR correction. 160 out of 196 path-non-random association with known biological processes and confirming statistical relevance of SHAP-prioritized pathways, Fig. 9.

**Fig. 9:**
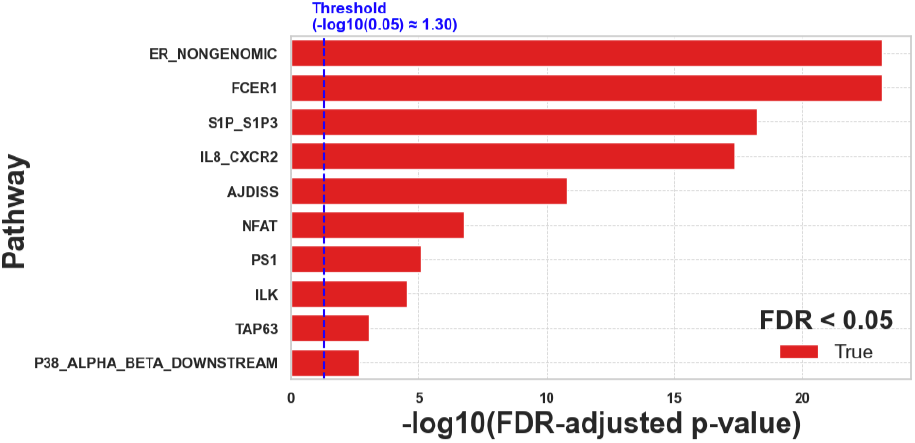
FDR-adjusted p-values of top ten enriched pathways. ways were significantly enriched (FDR < 0.05), indicating

Additionally, we pulled disgenet database [40] for literature validation. We found several evidences of the top ten significant genes across multiple tumor types. For instance, SRC is implicated in colon, breast, and bladder cancers, with known roles in tumor progression and EGFR/MAPK signaling activation (PMIDs: 9988270, 21357651, 19896475, 11723127).

PIK3R1 mutations contribute to oncogenic activation of the PI3K/AKT/mTOR pathway in endometrial and other cancers (PMID: 29636477). RHOA harbors somatic mutations in angioimmunoblastic T-cell lymphoma and diffuse-type gastric carcinoma, often in cooperation with TET2 mutations (PMIDs: 24413734, 24816255, 24413737). PIK3CA is among the most frequently mutated oncogenes across breast, colorectal, and ovarian cancers, with gain-of-function mutations driving tumorigenesis and showing therapeutic response to PI3K inhibitors such as alpelisib (PMIDs: 26266975, 15930273, 15520168, 37908459, 29899452).

## IV. Conclusion

We developed PathPCNet, a novel interpretable machine learning framework, integrating multi-omics data and biological pathways combined with PCA for drug response prediction. We applied SHAP to identify the significant pathways that influence the drug response. Additionally, we back-projected the SHAP scores to the original features (genes) using PCA loadings and ranked the genes based on their significance. Further, we provided the visual interpretation of SHAP scores for Morgan fingerprints of the drugs, which can help in designing better drugs. To the best of our knowledge, this is the first study to employ pathway-based principal component features for drug response prediction. Our framework can help with biomarker discoveries for drug response, as well as design better treatment strategies based on individual genetic profiles, as well as the molecular structure of the drugs.

A potential future work can be to integrate other omics data (e.g, DNA methylation) and the spatial data to study how other types of genomic data influence the tumor response. Since a tumor response is a complex biological phenomenon that is beyond a simple gene interaction, we believe integrating other omics data may provide better insights into the underlying omics profile that influences the drug response.

